# Structural Connectivity Correlates of Response to Electroconvulsive Therapy in Treatment-Resistant Depression

**DOI:** 10.1101/2025.03.18.643974

**Authors:** María Eugenia Samman, Leticia Fiorentini, María Lucía Fazzito, Delfina Lahitou Herlyn, Aki Tsuchiyagaito, Mariana N. Castro, Elsa Costanzo, Joan A. Camprodon, Cecilia Forcato, Salvador M. Guinjoan, Mirta F. Villarreal

## Abstract

**Background:** Electroconvulsive therapy (ECT) is the most effective option for treatment resistant depression (TRD). In this study, we sought to explore if structural connectivity of limbic networks has an association with response to ECT.

**Methods:** We studied 23 patients with TRD who underwent a course of bifrontal ECT, employing probabilistic tractography at baseline to assess structural connectivity between the thalamus (THA), posterior (PCC), subgenual cingulate cortices, anterior insula (aINS), amygdala and orbitofrontal and ventrolateral prefrontal cortices, hypothesizing that these hubs participate in the formation and refractoriness of depression symptoms. We also include 21 healthy subject as controls group (HC).

**Results:** Connectivity between left THA and left PCC was related to both baseline depression severity (R=0.504; p= 0.017) and clinical response (R=0.452; p=0.004). Right aINS-prefrontal connectivity was associated with less clinical response. Structural connectivity was globally higher in patients than in HC (F=2.488; p=0.007).

**Conclusions:** The association of ECT response with stronger structural connectivity between hubs supporting self-referential bodily experience as well as autobiographical memory encoding and retrieval deserves exploration as a predictor in persons with TRD. In turn, the right aINS is a major hub for the salience network and is involved in repetitive negative mentation. Stronger structural connectivity of this region may be a heuristically valid biomarker for refractory TRD. We discuss the potential of the present findings for the design of anatomically precise neuromodulation interventions that would be useful to treat TRD while circumventing cognitive side effects of ECT.

## 1. Introduction

Resistance to first-line treatments for major depressive disorder (MDD), including antidepressants and psychotherapies, is a common clinical problem, affecting about one third of all patients (Nemeroff, 2020). In this group, electroconvulsive therapy (ECT) remains an effective treatment option, reaching high response and remission rates (Espinoza and Kellner, 2022; Geddes et al., 2003). The mechanism of action and neurobiological correlates associated to response to ECT are currently largely unsettled, despite this technique being part of the therapeutic armamentarium for almost one century (Geddes et al., 2003).

Diverse studies employing functional connectivity have consistently implicated changes in default-mode network (DMN) connectivity as reflective of clinical response to ECT (Denier et al., 2023; Ten Doesschate et al., 2023; Verdijk et al., 2024). A recent multi-center cohort study explored the effective resting state connectivity correlates of severe depression and its treatment with ECT, including regions of three networks: DMN, salience network (SN), and central executive network (CEN). This study reported interactions particularly between the posterior cingulate cortex (in DMN), the insula (in SN), and anterior prefrontal regions (in CEN), associated with positive or negative treatment outcome (Ten Doesschate et al., 2023). Another recent, large multicenter study employing machine learning to define biomarkers of ECT response in depression, has implicated combined structural gray matter and functional connectivity features, revealing predictors of response also in midline structures like the precuneus, along with the thalamus (Bruin et al., 2024). In turn, burgeoning evidence by our group and others underscores the role of thalamo-cortical connectivity in depressive symptom formation, especially regarding self-referential negative mentation, a prominent feature of treatment-resistant MDD (Tsuchiyagaito et al., 2023; Yang et al., 2023).

On the other hand, there is a paucity of studies of anatomical disposition of white matter tracts subserving clinical improvement after ECT. The appropriate characterization of white matter connectivity correlates of ECT response would be desirable because it would open the possibility of selectively targeting edges that participate of specific large-scale circuits, thus ideally avoiding ECT adverse effects that result from the relative lack of specificity of this method. In fact, there is growing consensus that new developments in circuit neuromodulation need to focus on white matter connections, rather than grey matter circuit hubs, and therefore an understanding of structural connectivity patterns associated to ECT response would have significant clinical heuristic value (Coenen et al., 2019; Chan et al., 2024; Figee et al., 2022; Sanchez et al., 2023). Tsolaki et al. (2021) recently put to test the hypothesis that response to ECT is dependent on the structural connectivity of the subgenual cingulate cortex (SGCC), as could be predicted from the prominent role of this region in the production of depressive symptoms and their response to a variety of treatments. They demonstrated that structural connectivity between the SGCC with the rest of the cingulate gyrus and with medial prefrontal cortex structures was lower in responders compared to non-responders. Intriguingly, non-responders were similar to healthy controls in their pattern of connectivity, thus suggesting that SGCC structural connectivity may work as a robust biomarker of response to ECT, rather than a neurobiological signature of depression.

In the present study, we sought to expand prior evidence on neurobiological signatures of response to ECT by incorporating hypotheses generated from recent studies assigning a critical role of connectivity on the pathophysiology of depressive symptoms and treatment response. Specifically we explored: 1- posterior cingulate cortex (PCC) as a major node of the DMN; 2- anterior insula (aINS) as the main node of the SN (Dunlop et al., 2019; Fonseka et al., 2018); 3- orbitofrontal cortex given its critical role in both affective processing and reward (Huang et al., 2023; Rolls et al., 2020); 4- SGCC given its pivotal role in depressive symptom development, and as a major target for effective neuromodulation treatments (Cattarinussi et al., 2022; Sobstyl et al., 2022) and its connectivity with ventrolateral prefrontal cortex (VLPFC), and 5- subcortical structures including the thalamus (Brown et al., 2017; Hwang et al., 2022) and amygdala (AMY), whose volume was reported to be a predictor of response to ECT (Dunlop et al., 2019; Fonseka et al., 2018). Networks involving most of the mentioned regions (OFC, AMY, VLPFC, PCC) were particularly found to predict response to repetitive transcranial magnetic stimulation trough strongest resting state connectivity (Drysdale et al., 2017), although the evidence of abnormalities in the functional or structural connectivity of most of these regions and their potential as biomarkers of successful response appears to depend on the type of treatment (Tura and Goya-Maldonado, 2023). Based upon prior studies we specifically hypothesized that lower structural connectivity between the SGCC, anterior insula and thalamus would be a marker of response to ECT, along with greater connectivity between posterior DMN components and subcortical regions. With this aim, we studied 23 consecutive patients with treatment-resistant depression (TRD) who underwent a full course of bilateral ECT.

## 2. Materials and Methods

### 2.1. Participants

Twenty-nine consecutive patients with TRD eligible to receive ECT, were recruited from the Service of Psychiatry of the Fleni Foundation, Buenos Aires. Additionally, twenty one healthy control subjects (HC) without history of psychiatric disorders were also included (Table 1) to establish normative values and test for overall diagnostic group differences. All participants provided written informed consent as authorized by the local bioethics committee at Fleni. The Supplementary Material shows the inclusion and exclusion criteria for the study.

**Table 1.** Demographic characteristics and ECT Response.

|  | TRD<br>(n=23) | HC<br>(n=20) | Statistics | P |
| --- | --- | --- | --- | --- |
| Gender (male/female) | 13/10 | 9/11 | $\chi^2 = 0.201$ | 0.654 |
| Age (years) | 49.3 ± 14.5 | 28.7 ± 5.9 | W= 417.5 | < 0.05 |
| Education (years) | 13.7 ± 3.4 | 16.1 ± 2.5 | W= 149 | 0.047 |
| HAM-D Baseline | 24.9 ± 7.5 | 2.4 ± 2.4 | W= 460 | < 0.05 |
| Age of onset (years) | 34.49 ± 15.1 |  |  |  |
| Duration of illness (months) | 178.7 ± 107.9 |  |  |  |
| Duration of episode (months) | 17.6 ± 15.5 |  |  |  |
| Antidepressants #/n | 21/23 |  |  |  |
| Antipsychotic #/n | 20/23 |  |  |  |
| **Responder #/n** | 15/23 |  |  |  |
| Post ECT HAM-D | 4.7 ± 3.1 |  |  |  |
| HAM-D % | 92.3 ± 13.3 |  |  |  |
| **Non Responder #/n** | 8/23 |  |  |  |
| Post ECT HAM-D | 17.4 ± 7.0 |  |  |  |
| HAM-D % | 10.5 ± 17.1 |  |  |  |

Values shown as mean ± SD. HAM-D: Hamilton Depression Rating Scale. W: Mann Whitney test statistic

Patients underwent 12 session of ECT as detail in Supplement. The Hamilton Depression Rating Scale-17 items (HAM-D; Hamilton, 1960) was used to assess the symptom severity of the patient group before and after the ECT. The HC group also underwent assessment using the HAM-D. Patients whose HAM-D score decreased by at least 50% after treatment were considered responders (R) (Nierenberg and DeCecco, 2001), while those who did not reach the 50% were considered non-responders (NR). For further analysis, we define the change in HAM-D (*HAM-D%)* as:

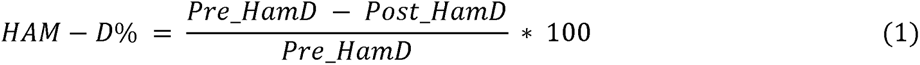

where proportional, rather than absolute, symptom changes were employed to mitigate an overestimation of the therapeutic response to ECT as a function of higher HAM-D scores (Table 1).

Patients were scanned on a 3T GE Discovery 750 MRI system (Fleni Foundation) within one week prior to the first ECT session. HC were also scanned with the same system and parameters (Supplement). Diffusion images were first pre-process to correct current and movement distortions, and posteriorly processed to build up the distributions on diffusion parameters per voxel (see Supplement).

### 2.2. Probabilistic tractography

Probabilistic tractography was run at baseline using seed masks manually built on seven bilateral regions of interest (ROIs). Each seed region was conformed as a sphere of 5 mm radius centered in the MNI coordinates described in Table 2. This selection was hypothesis driven, and the location was based on extant literature assigning a preeminent role in the development of depressive symptoms, their resistance, and their response to treatment, to a series of cortical and subcortical grey matter hubs (Koenigs et al., 2008; Runia et al., 2022; Sobstyl et al., 2022).

**Table 2.** Regions of interest (ROI) and seed placement.

| **Region name**<br>**(Right and Left)** | **Abbreviation**<br>**(R./L.)** | **MNI coordinates**<br>**(+) Right, (-) Left** |
| --- | --- | --- |
| Amygdala | AMY | (± 23; -5; -19) |
| Anterior insula | aINS | (± 36; 13; 5) |
| Orbito frontal cortex | OFC | (± 16, 42, -18) |
| Posterior cingulate cortex | PCC | (± 5; - 50; 30) |
| Posterior ventrolateral prefrontal cortex | pVLPFC | (± 51; 20; 5) |
| Subcallosal cingulate gyrus | SGCC | (± 6; 25; -9) |
| Thalamus | THA | (± 10; -21; 10) |

Connectivity values between ROIs were evaluated as described in Supplement. Finally, the probability of each pair was calculated as the mean connectivity value between pairs *i*- *j* and *j*- *i* normalized to each individual’s maximum connectivity value as follow:

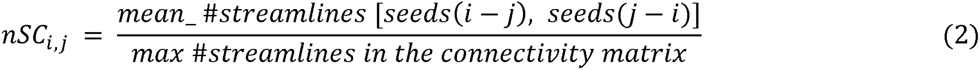

where n*SC* is the normalized structural connectivity, and *i* and *j* account for the seed masks (Table 2). Therefore, the range for all scores are between 0-1.

### 2.3. Statistical analysis

First, to explore the relationship between the initial depressive symptomatology and structural connectivity, we correlated the HAM-D score at baseline with the *nSC_i,j_*. The same analysis was also performed for HC group. Second, to explore the relationship between the response to treatment and the structural connectivity for patients, we correlate the HAM-D % with the *nSC_i,j_* values. In all cases, we used Spearman correlations methods, and included age as nuisance covariate. Additionally, to check for potential confounds on the correlations with the HAM-D% change, due to their initial values in the HAM-D, we also assessed the *nSC* vs HAM-D% correlation by adding the baseline HAM-D score as a nuisance covariate. As this study represents an exploratory approach, we show results with statistical significance from p ≥ 0.05, without multiple comparison corrections to minimize false negative results.

Finally, to explore differences in the strength of connectivity between patients biotypes, based on response type to ECT, and control subjects, a general linear model was implemented on those pairs of ROIS that demonstrate a significant correlation with response to ECT. Age and HAM-D at baseline were included as nuisance covariates.

## 3. Results

### 3.1. Demographics and Clinical Response

Table 1 shows the demographic and clinical characteristics of the sample. After ECT, the reduction in HAM-D score was 21.7 ± 7.9 in responders (n = 15) and 5.0 ± 4.2 in non-responders (n = 8). Patients and HC did not differ significantly in gender, although patients were older than control subjects (W= 437.5, p < 0.05), and have fewer years of education (W = 149, p = 0.047).

### 3.2. Correlation Analyses

Eleven structural connections were correlated with baseline HAM-D in patients with TRD, as shown in Figure 1, panel A and Table 3. The connections between THA and PCC were positively correlated with the baseline HAM-D score. In contrast, the remaining connections exhibited the opposite pattern, with higher n*SC* values corresponding to lower clinical severity. Figure 1, panel B top and middle, shows the relationship between the structural connectivity of two relevant brain connections (left PCC – right THA and left aINS – left SGCC, respectively) and the baseline Ham-D score for patients, and for HC (bottom panel). HC did not present the associations between HAM-D and structural connections found in TRD. Conversely, they showed significant inverse relationships for connections between different brain areas, as shown in Table 3.

**Figure 1:**
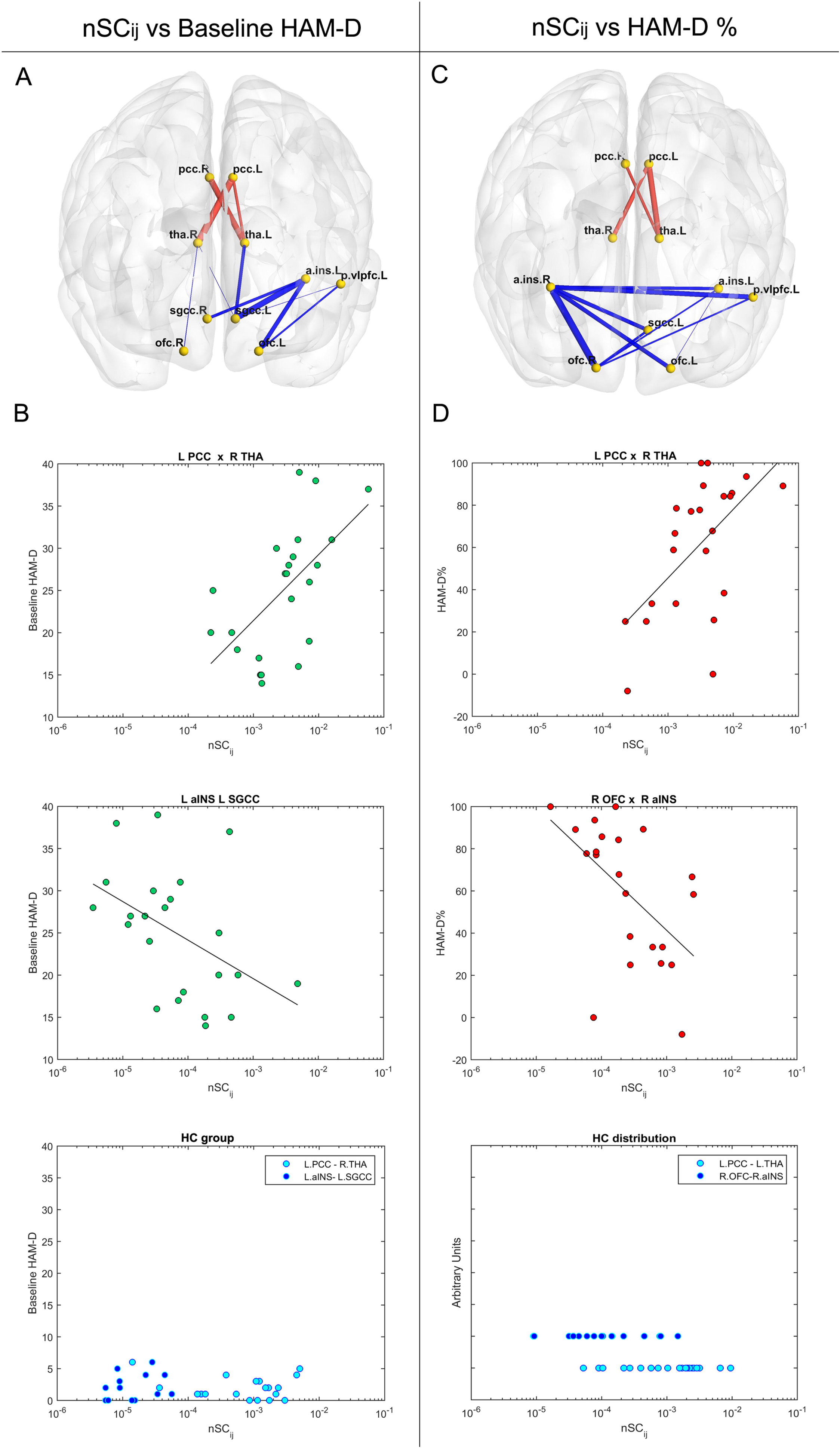
Structural connectivity and baseline Ham-D (left column), and response to ECT (right column). Panels A and C show the significant connections on brain templates for baseline Ham-D (A), and Ham-D % (C), with the links representing the Spearman R values (red positive, blue negative). Panel B top and middle show scatter plots of the relationship between the normalized structural connectivity (n*SC*) of two connections and baseline Ham-D in patients. Panel B bottom shows the relationship of the same connections in HC (symbols cyan and blue for each one). Panel D top and middle show scatter plots of the relationship of the n*SC* in two connections and the clinical response Ham-D % for patients. Panel B bottom shows the n*SC* distribution of HC in the same connections, plotted with an arbitrary y-axis (symbols cyan and blue for each one). L: left, R: right, aINS: anterior insula, OFC: orbitofrontal cortex, PCC: posterior cingulate cortex, THA: thalamus, SGCC: subgenual cingulate cortex.

**Table 3:**
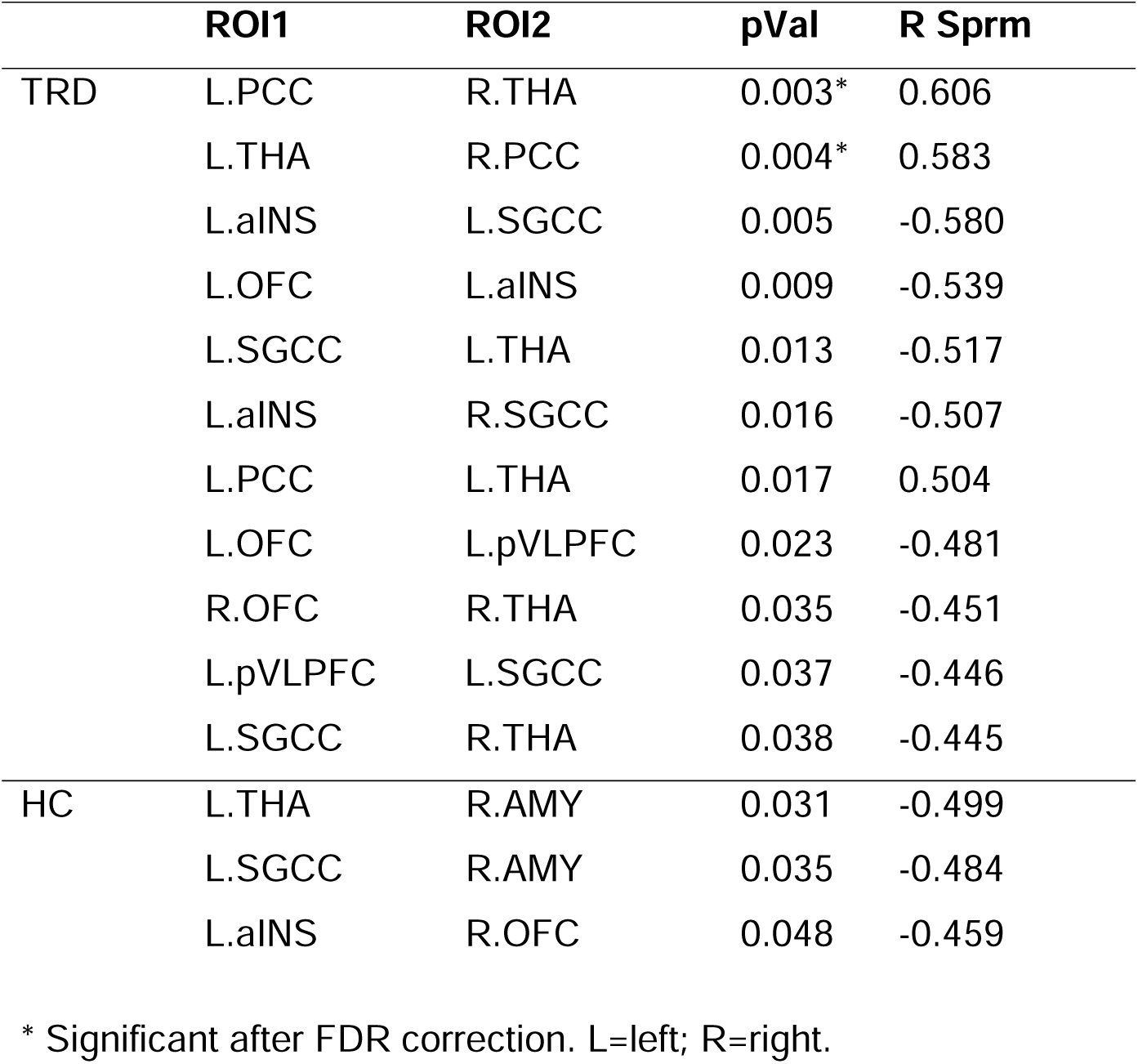
Correlations between structural connectivity and depression symptoms.

Changes in HAM-D score after ECT treatment have significant associations with structural connectivity for 12 pairs of brain regions as shown in Figure 1, panel C and Table 4. The strength of connectivity between bilateral THA with left PCC, and left THA with right PCC has a direct relationship with HAM-D% changes. Instead, the remainder connections had a negative correlation with response to treatment. Figure 1, panel D top and middle, shows the relationship between the structural connectivity of two relevant pairs (left PCC–THA and right OFC–aINS, respectively) and the HAM-D% for patients. Figure 1D, bottom panel, shows the distribution of n*SC* in the HC sample in those same pairs.

**Table 4:** Correlations between structural connectivity and change in depression symptoms.

| **ROI1** | **ROI2** | **pVal** | **R Sprm** |
| --- | --- | --- | --- |
| R.OFC | R.aINS | 0.003* | -0.598 |
| L.PCC | L.THA | 0.006* | 0.566 |
| L.pVLPFC | R.aINS | 0.009* | -0.544 |
| L.PCC | R.THA | 0.011* | 0.533 |
| L.OFC | R.aINS | 0.011* | -0.527 |
| L.SGCC | R.aINS | 0.013* | -0.521 |
| L.aINS | R.aINS | 0.017 | -0.502 |
| L.THA | R.PCC | 0.020 | 0.492 |
| L.SGCC | R.OFC | 0.025 | -0.477 |
| L.pVLPFC | R.OFC | 0.030 | -0.462 |
| L.aINS | R.OFC | 0.033 | -0.456 |
| L.OFC | L.aINS | 0.048 | -0.427 |

\* Significant after FDR correction. L=left; R=right.

Both panels A and C of Figure 1, depict that thalamic-PCC connections and the left aINS–OFC correlates with both baseline HAM-D and HAM-D%. However, when the HAM-D value at baseline was added as a nuisance covariate in the correlation analysis only left THA–PCC connection remained significant. Figure 2 and Table 5 summarize all structural connectivity with significant values after controlling for HAM-D.

**Figure 2:**
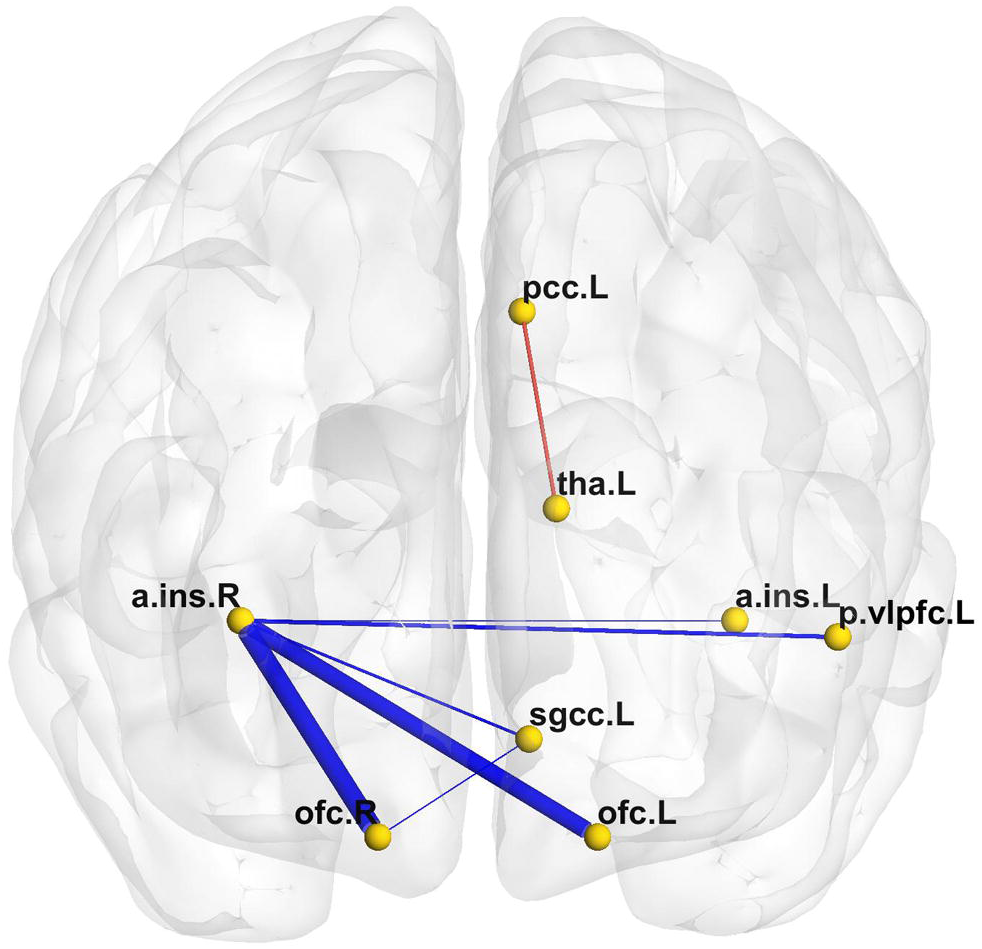
Structural connectivity and response to ECT treatment with baseline HAM-D score as covariate. The significant connections associated with HAM-D% are shown on a brain template using red links for positive correlations and blue links for the negative correlations.

**Table 5:** Correlations between structural connectivity and change in depression symptoms adding HAM-D as nuisance covariate.

| **ROI1** | **ROI2** | **pVal** | **R Sprm** |
| --- | --- | --- | --- |
| R.OFC | R.aINS | 0.009 | -0.555 |
| L.OFC | R.aINS | 0.012 | -0.541 |
| L.SGCC | R.aINS | 0.024 | -0.491 |
| L.pVLPFC | R.aINS | 0.034 | -0.465 |
| L.PCC | L.THA | 0.004 | 0.452 |
| L.aINS | R.aINS | 0.047 | -0.437 |
| L.SGCC | R.OFC | 0.048 | -0.433 |

### 3.3. Structural connectivity intergroup differences

Five structural connections showed a significant effect between the three groups (R, NR and HC, see Table 5 and Figure 3). NR group exhibited higher scores of n*SC* compared to HC group in the connections involving the R.aINS with the L.OFC, the L.pVlPFC and the L.aINS (p = 0.012, p = 0.009, p = 0.014, respectively), indicated with one asterisk in the plots of Figure 3 B,C,E. The same connections was significant higher for NR compared to R group (p = 3.10^-5^, p = 0.024, p = 0.032 respectively), with the addition of the L.SGCC–R.OFC (p =0.007) indicated with two asterisk in the plots of Figure 3 B,C,E,F. The pair R.OFC–R.aINS (Figure 3 A), only show a tendency for NR > HC (p = 0.054). Instead, R group had significant greater connectivity than NR in the L.PCC–L.THA connection (p = 0.009), indicated with three asterisk in Figure 3 D.

**Figure 3:**
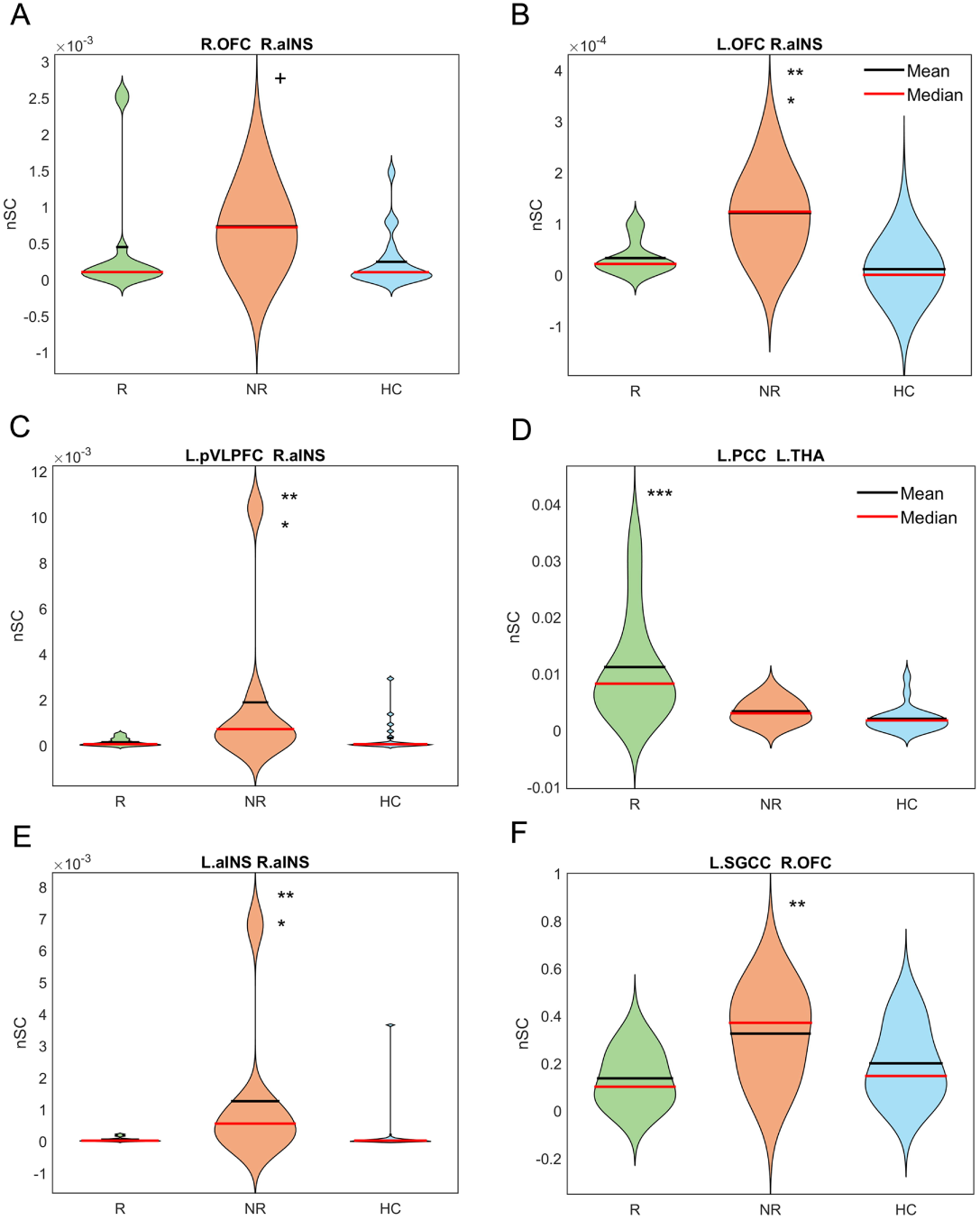
Structural connectivity intergroup differences. The violin plots show the distribution of *nSC* in the three groups: R for responders (green), NR for non-responders (orange) and HC for healthy controls (cyan) for the connections with significant group effect. Asterisks beside a distribution indicate a higher n*SC* score than (*) HC, (**) R, (***) NR. Plus symbol (+) indicate a tendency of NR >HC. L: left, R: right, aINS: anterior insula, OFC: orbitofrontal cortex, pVLPFC: posterior ventrolateral prefrontal cortex, PCC: posterior cingulate cortex, THA: thalamus, SGCC: subgenual cingulate cortex.

Figure 4 depicts individual examples of tractography connecting the L.PCC–L.THA pair, where a R exemplary participant shows higher values of n*SC* than NR (panels A for R, and B for NR participants). Figure 4 second line (panels C for R, and D for NR participants), displays the tractography connecting the L.OFC–R.aINS pair for the same individuals. In this case, the amount of n*SC* is higher for NR with respect to R subjects. The selection criterion for both patients was a similar Ham-D at baseline (27 for R, and 25 for NR patients).

**Figure 4.**
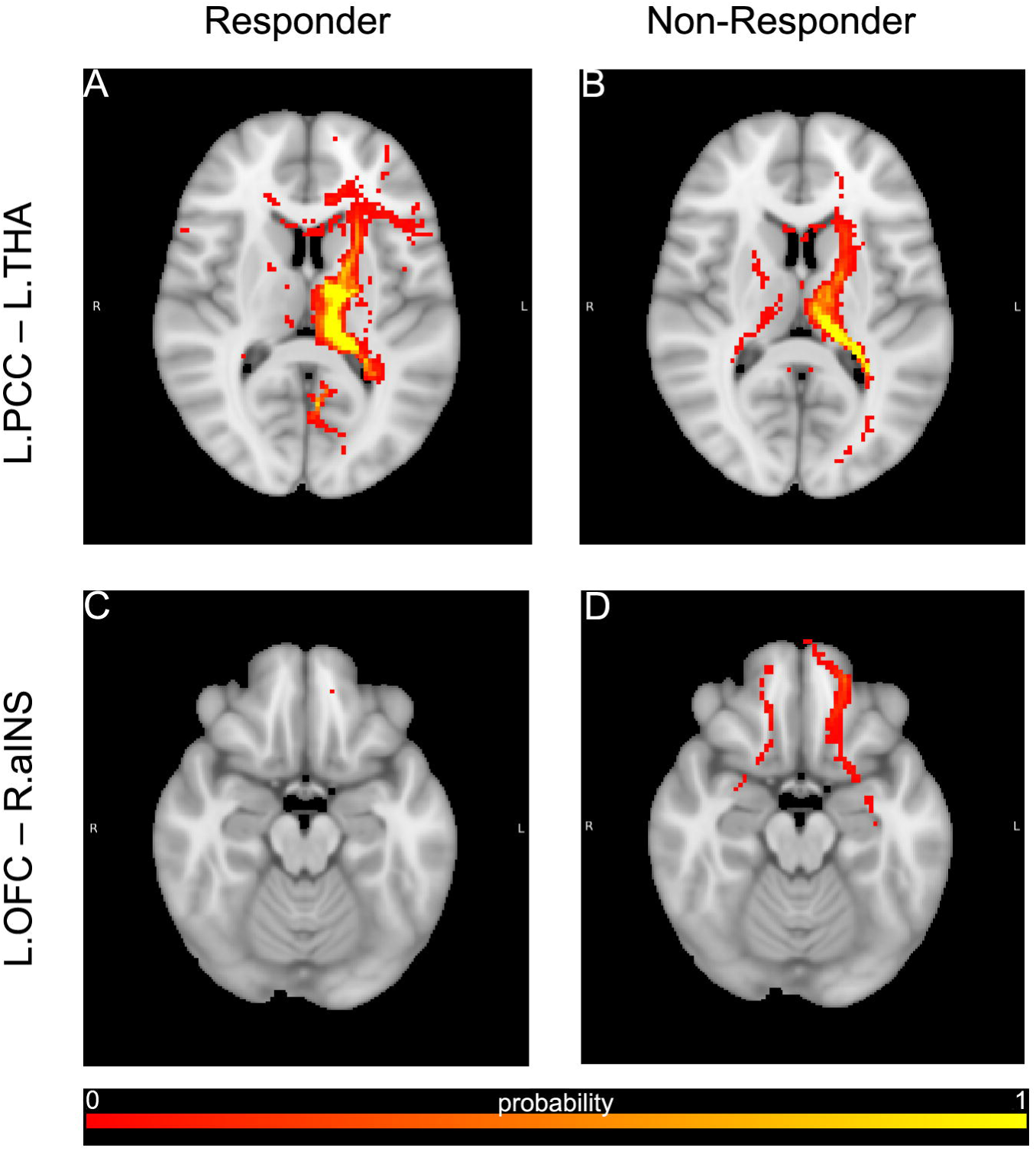
Individual examples of tractographies. A patient who responded (HAM-D at baseline = 27, HAM-D% = 100) is shown in Panels A and C (first column). A patient who did not respond to treatment (HAM-D at baseline = 24, HAM-D% = -8) is shown in Panels B and D (second column). Connection L.PCC—L.THAL is show in the first line (A and B), and L.OFC—R.aINS in the second line (C, D).

## 4. Discussion

The present study explored the relationship of structural connectivity between a series of gray matter hubs which have been shown in extant literature to have a role in the development of MDD symptoms, their resistance to treatment, and clinical response to a full course of ECT. The main findings of the study include: 1) the strength of structural connections between the left THA and the left PCC is associated with both more intense pre-treatment depressive symptomatology and greater likelihood of response to ECT, 2) the strength of structural connections between the right aINS and bilateral OFC, left pVLPFC, left SGCC and left aINS, as well the right OFC – left SGCC connections is associated with a worse response to ECT, 3) individuals who responded to treatment exhibited higher *nSC* scores in the left THA–PCC connection compared to non-responders, 4) non-responder patients exhibit higher connectivity than HC between the right aINS and left OFC, left pVLPFC, and left aINS, 5) non-responder patients exhibit higher connectivity than responder patients between the right aINS and left OFC, left pVLPFC, left aINS, and between the right OFC and left SGCC, and 6) HC show a negative correlation between their depressive symptom score and left THA– right AMY, left SGCC– right AMY and left aINS– right OFC, with none of these connections overlapping with those seen in patients.

Our study provides evidence that greater proportional connectivity in the left THA – PCC connection in the responder group might facilitate response to treatment. In contrast, in most of the structures involving the aINS and the prefrontal ROIs, where the correlation was inverse to clinical response, non-responder patients exhibit higher connectivity than both HC and responders, suggesting that stronger connectivity may hinder a favorable response to treatment.

As predicted, structural connectivity between subcortical regions, specifically the thalamus, and a major node of the DMN, namely the PCC, was strongly related to response to ECT. Further, this variable was the only structural connectivity pattern associated to greater depression severity prior to treatment. With respect to the SGCC and aINS structures, we found that lower connectivity with prefrontal regions and not subcortical are linked to a greater response, as initially hypothesized. If replicated, these observations can shed light on the pathophysiology of treatment resistant depression and the mechanisms of response to treatment to ECT in this group.

### 4.1. Thalamo-Posterior Cingulate Cortex Structural Connectivity was Associated with Response to ECT

Our finding of greater structural connectivity between the thalamus and the PCC as both a correlate of depression severity and of likelihood of response to ECT can be best interpreted in light of the networks in which the PCC participates. The PCC is a grey matter hub involved in the regulation of emotion and memory, and executive control. The PCC establishes reciprocal connections with diverse nuclei within the thalamus, reflecting its nodal position in self-referential phenomena including mentation, body localization, and episodic or autobiographical memory (Foster et al., 2023; Leech and Sharp, 2014; Rolls, 2019). In this setting, conceivably a greater structural connectivity might subserve persistent and increased negative self-referential mentation, which is characteristic of depression. In fact, we have previously posited that a major potential mechanism of treatment-resistant in depression is a persistent cycle of retrieval and reconsolidation (including REM sleep consolidation) of negatively laden emotional autobiographical memories, manifested clinically as rumination, or facilitated by this symptom (Guinjoan et al., 2021). Indeed, both ECT and sleep deprivation could interfere with this cognitive phenomenon (Guinjoan et al., 2021). Compatible with this interpretation and our present results, ECT has been recently observed to interfere with functional connectivity of posterior DMN hubs including the PCC (Gbyl et al., 2024). That the large-scale structural circuit involving the PCC shows specificity for connections with the thalamus lends support to the importance of this structure as a major gateway conveying information from subcortical structures, and thus with a major role in the formation of depressive symptoms in general, as recently proposed (Yang et al., 2023). Thus, it has been suggested that excessive bottom-up negative affect sensory information, along with deficit top-down suppression of negative emotions may both be critical for the development of depression, and the thalamus occupies a central role in the production of symptoms and their resolution via specific disruption of its connections, especially those that enter the cingulum bundle and reach its different components (Davidson et al., 2022; Velasco et al., 2005). Considering this evidence, our finding of white matter connections linking thalamus and PCC as related to both depression severity and its improvement with ECT, suggests that these well-characterized and easily imaginable white matter tracts deserve further exploration as a specific target for anatomically precise deep-brain neuromodulation methods, including innovative tools like focused ultrasound (Philip and Arulpragasam, 2023).

### 4.2. Right Anterior Insula Structural Connectivity was Associated with Non-Response to ECT

We found higher connection between the right anterior insula and prefrontal regions in non-responders compared to HC and responder patients, which inversely correlates with clinical improvement. The right anterior insula (R.aINS) is a crucial hub within the salience network involved in various cognitive and emotional processes. Its functions extend across multiple domains, particularly in the context of salience detection and interoception (Craig, 2009; Haruki and Ogawa, 2021; Menon and Uddin, 2010). Interoceptive awareness encompasses the perception of internal bodily states such as hunger, pain, or emotional discomfort. This function is essential for emotional experiences and self-awareness, allowing individuals to relate to their own feelings and those of others (Uddin et al., 2017). In that context, the RaINS plays a crucial role in the pathophysiology of depression.

Diverse studies have shown that individuals with MDD exhibit significant alterations in the functional and effective connectivity of the R.aINS. For instance, it was reported that patients with depression had reduced connectivity from the R.aINS to the dorsolateral prefrontal cortex, which is essential for cognitive control and emotional regulation, and additionally, increased connectivity between the amygdala and RAI, suggesting a heightened emotional response to negative stimuli (Kandilarova et al., 2018). This altered connectivity is associated with the severity of depressive symptoms, indicating that disruptions in these neural circuits may contribute to the experience of depression. Increased activation of the R.aINS has also been linked to greater rumination in depressed patients, suggesting that this region may facilitate a persistent focus on negative emotional states. This hyperactivation can lead to a bias toward processing negative information more intensely than neutral or positive stimuli (Sliz and Hayley, 2012). Furthermore, disruptions in top-down emotional regulation involving the R.aINS can exacerbate feelings of sadness and negative affect in individuals with depression (Dobrushina et al., 2021; Sliz and Hayley, 2012). Structurally, the R.aINS is interconnected with various brain regions in the frontal lobes through white matter tracts. These include connections to the OFC, the VLPFC and other areas involved in emotional and cognitive processing (Zhao et al., 2023). Hence, considering the mention findings and characteristics, the augmentation of fiber bundles connecting the R.aINS with other prefrontal regions in non-responder patients, besides a negative contribution with the clinical improvement are in agreement with the scenario of overwhelming self-awareness, rumination, and lack of negative emotions regulation.

### 4.3. Structural Connectivity Associations in HC

Our analysis revealed stronger connectivity between the right AMY and both the left SGCC and the left THA, as well as between the right OFC and left aINS with lower HAM-D scores in HC. Or put in another words, the weaker the connections, the higher the scores in the depression scale. The AMY, together with the OFC and the aINS, are involved in the processing of salient stimuli and have been considered central hubs within the affective salience network (Damborská et al., 2020; Pessoa and Adolphs, 2010; Rolls et al., 2020). The limbic-thalamocortical circuit also plays an important role in the regulation of emotional responses, a process that is altered in various psychiatric disorders, particularly in MDD. The integrity of the different tracts connecting these regions (forceps minor, anterior thalamic radiations, uncinate fasciculus and cingulate bundle) has been shown to be reduced in the pathophysiology of depression, leading to emotional imbalance and the predominance of negative emotions (Bracht et al., 2015; Lewinn et al., 2014; Mincic, 2015). In this context, our results in HC are consistent with this line, showing that a more robust connectivity of these regions, linked to a better integrity of the tracts involved, is associated with mental health.

## 5. Limitations

Contrary to our hypothesis, we did not observe a relationship between the structural connectivity between thalamus and SGCC, in either depression severity or likelihood of response to ECT. In a landmark communication, Tsolaki and coworkers (2021) observed that ECT responders had a lower structural connectivity between the SGCC and the medial prefrontal cortex, compared to both non-responders and HC. The latter observation suggests that this finding represented a biomarker of treatment response, unrelated to depression pathophysiology, given that patients with depression who did not respond to ECT, and whose severity was comparable to that of responders, did not differ in this regard from HC. One of the possible reasons for the discrepancy between this study and our results, is that we quantified streamlines connecting SGCC with similar but not identical areas of the cortex and subcortical regions. Therefore, our negative results may complement, rather than contradict, this previous observation.

Our work had diverse limitations that need to be considered in the interpretation of results.

First, the small sample size could have resulted in type II errors, including our negative observation on the relationship between structural connectivity of the thalamus and the SGCC, and symptom improvement after ECT. Second, the study was performed in patients from a single institution, thus limiting the generalizability of the present results. Third, the control group was significantly younger than patients, and although we controlled for age effects in the statistical model, the white matter structure is known to change with age; nevertheless, we found significant differences between patients group also, who did not differed in age (p = 0.95, not shown previously). Fourthly, patients have fewer years of education compared to HC, and while this could affect myelination, its impact is more significant during the early years of development than in the later university years. In this context, the age difference is more relevant in our sample. Finally, although we used a standardized diffusion tensor probabilistic analysis pipeline, other methods could have resulted in different connectivity observations.

## 6. Conclusion

The strength of specific left thalamo-left PCC structural connections at baseline correlates with both symptomatology before the treatment and clinical outcome of ECT treatment, with significantly stronger connectivity for responding patients. The result suggests that this finding represented a biomarker of treatment response, unrelated to depression pathophysiology. For the other side, right anterior insula stands out as an important hub whose links with bilateral OFC, left pVLPFC, left SGCC and left aINS correlates only with clinical outcome, but in an inverse relationship, i.e., the greater the strength, the worse the outcome. Consistent with this observation, non-responders have significantly more connections between most of the mentioned nodes than responders and HC. For that reason, the strength of those connections may conceivably help differentiate disease biotypes in TRD population of patients, specifically between responders and non-responders to ECT. In consequence, these connectivity patterns may serve as potential clinical biomarkers for ECT response and help in treatment selection. Most important, the definition of structural correlates of response to ECT warrants their exploration as a target for anatomically precise, deep neuromodulation tools, including traditional surgical procedures, and recently developed non-invasive devices (Philip and Arulpragasam, 2023).

## Supporting information

Supplementary Material_ Samman et al 2025

## 7. CRediT authorship contribution statement

SMG and MFV developed the scientific project. EC, MLF, DLH and LF recruit the participants and performed the clinical and neuropsychological evaluations. MLF and MFV obtained MR acquisitions. EC and MLF performed the ECT sessions. MES and MFV analyzed the diffusion data. MES, MFV and SMG wrote the first draft of the manuscript. MNC, CF, JAC and AT contributed in the final version of the manuscript. All authors critically reviewed the manuscript and have approved the final manuscript.

## 10. Acknowledgements

We are grateful to the patients for agreeing to participate in the study. MES is a graduate student in the Environment and Health Applied Sciences Doctoral Program (DCAAS) at UNICEN, Argentina.

## References

Bracht, T., Linden, D., Keedwell, P., 2015. A review of white matter microstructure alterations of pathways of the reward circuit in depression. J. Affect. Disord. 187, 45–53. 10.1016/j.jad.2015.06.041.

Brown, E.C., Clark, D.L., Hassel, S., MacQueen, G., Ramasubbu, R., 2017. Thalamocortical connectivity in major depressive disorder. J. Affect. Disord. 217, 125–131. 10.1016/j.jad.2017.04.004.

Bruin, W.B., Oltedal, L., Bartsch, H., Abbott, C., Argyelan, M., Barbour, T., Camprodon, J., Chowdhury, S., Espinoza, R., Mulders, P., Narr, K., Oudega, M., Rhebergen, D., Ten Doesschate, F., Tendolkar, I., van Eijndhoven, P., van Exel, E., van Verseveld, M., Wade, B., van Waarde, J., Zhutovsky, P., Dols, A., van Wingen, G., 2024. >Development and validation of a multimodal neuroimaging biomarker for electroconvulsive therapy outcome in depression: a multicenter machine learning analysis. Psychol. Med. 54, 495–506. 10.1017/S0033291723002040.

Chan, J.L., Carpentier, A.V., Middlebrooks, E.H., Okun, M.S., Wong, J.K., 2024. Current perspectives on tractography-guided deep brain stimulation for the treatment of mood disorders. Expert Rev.

Coenen, V.A., Schlaepfer, T.E., Bewernick, B., Kilian, H., Kaller, C.P., Urbach, H., Li, M., Reisert, M., 2019. Frontal white matter architecture predicts efficacy of deep brain stimulation in major depression. Transl. Psychiatry. 9, 197. 10.1038/s41398-019-0540-4.

Craig, A.D., 2009. How do you feel--now? The anterior insula and human awareness. Nat. Rev. Neurosci. 10, 59–70. 10.1038/nrn2555.

Damborská, A., Honzírková, E., Barteček, R., Hořínková, J., Fedorová, S., Ondruš, Š., Michel, C.M., Rubega, M., 2020. Altered directed functional connectivity of the right amygdala in depression: high-density EEG study. Sci. Rep. 10, 4398. 10.1038/s41598-020-61264-z.

Davidson, B., Eapen-John, D., Mithani, K., Rabin, J.S., Meng, Y., Cao, X., Pople, C.B., Giacobbe, P., Hamani, C., Lipsman, N., 2022. Lesional psychiatric neurosurgery: meta-analysis of clinical outcomes using a transdiagnostic approach. J. Neurol. Neurosurg. Psychiatry 93, 207–215. 10.1136/jnnp-2020-325308.

Denier, N., Walther, S., Breit, S., Mertse, N., Federspiel, A., Meyer, A., Soravia, L.M., Wallimann, M., Wiest, R., Bracht, T., 2023. Electroconvulsive therapy induces remodeling of hippocampal co-activation with the default mode network in patients with depression. Neuroimage Clin. 38, 103404. 10.1016/j.nicl.2023.103404.

Dobrushina, O.R., Arina, G.A., Dobrynina, L.A., Novikova, E.S., Gubanova, M.V., Belopasova, A.V., Vorobeva, V.P., Suslina, A.D., Pechenkova, E.V., Perepelkina, O.S., Kremneva, E.I., Krotenkova, M.V., 2021. Sensory integration in interoception: Interplay between top-down and bottom-up processing. Cortex. 144, 185–197. 10.1016/j.cortex.2021.08.009.

Drysdale, A.T., Grosenick, L., Downar, J., Dunlop, K., Mansouri, F., Meng, Y., Fetcho, R.N., Zebley, B., Oathes, D.J., Etkin, A., Schatzberg, A.F., Sudheimer, K., Keller, J., Mayberg, H.S., Gunning, F.M., Alexopoulos, G.S., Fox, M.D., Pascual-Leone, A., Voss, H.U., Casey, B.J., Dubin, M.J., Liston, C., 2017. Resting-state connectivity biomarkers define neurophysiological subtypes of depression. Nat Med. 23(1), 28–38. 10.1038/nm.4246.

Dunlop, K., Talishinsky, A., Liston, C., 2019. Intrinsic Brain Network Biomarkers of Antidepressant Response: a Review. Curr Psychiatry Rep. 21(9), 87. 10.1007/s11920-019-1072-6.

Espinoza, R.T., Kellner, C.H., 2022. Electroconvulsive Therapy. N Engl J Med. 386(7), 667–672. 10.1056/NEJMra2034954.

Figee, M., Riva-Posse, P., Choi, K.S., Bederson, L., Mayberg, H.S., Kopell, B.H., 2022. Deep Brain Stimulation for Depression. Neurotherapeutics. 19(4), 1229–1245. 10.1007/s13311-022-01270-3.

Fonseka, T.M., MacQueen, G.M., Kennedy, S.H., 2018. Neuroimaging biomarkers as predictors of treatment outcome in Major Depressive Disorder. J. Affect Disord. 233, 21–35. 10.1016/j.jad.2017.10.049.

Foster, B.L., Koslov, S.R., Aponik-Gremillion, L., Monko, M.E., Hayden, B.Y., Heilbronner, S.R., 2023. A tripartite view of the posterior cingulate cortex. Nat. Rev. Neurosci. 24(3), 173–189. 10.1038/s41583-022-00661-x.

Gbyl, K., Labanauskas, V., Lundsgaard, C.C., Mathiassen, A., Ryszczuk, A., Siebner, H.R., et al., 2024. Electroconvulsive therapy disrupts functional connectivity between hippocampus and posterior default mode network. Prog. Neuropsychopharmacol. Biol. Psychiatry 132, 110981. 10.1016/j.pnpbp.2023.110981.

Geddes, J., Carney, S., Cowen, P., Goodwin, G., Rogers, R., Dearness, K., Tomlin, A., Eastaugh, J., Freemantle, N., Lester, H., Harvey, A., Scott, A., 2003. Efficacy and safety of electroconvulsive therapy in depressive disorders: A systematic review and meta-analysis. The Lancet 361, 799–808. 10.1016/S0140-6736(03)12705-5.

Guinjoan, S.M., Bär, K.J., Camprodon, J.A., 2021. Cognitive effects of rapid-acting treatments for resistant depression: Just adverse, or contributing to clinical efficacy? J Psychiatr Res 140, 512–521. 10.1016/j.jpsychires.2021.06.026.

Hamilton, M., 1960. A rating scale for depression. J. Neurol. Neurosurg. Psychiatry. 23, 56–62. 10.1136/jnnp.23.1.56.

Haruki, Y., Ogawa, K., 2021. Role of anatomical insular subdivisions in interoception: Interoceptive attention and accuracy have dissociable substrates. Eur. J. Neurosci. 53, 2669–2680. 10.1111/ejn.15157.

Huang, H., Rong, B., Chen, C., Wan, Q., Liu, Z., Zhou, Y., Wang, G., Wang, H., 2023. Common and distinct functional connectivity of the orbitofrontal cortex in depression and schizophrenia. Brain Sci. 13, 997. 10.3390/brainsci13070997.

Hwang, W.J., Kwak, Y.B., Cho, K.I.K., Lee, T.Y., Oh, H., Ha, M., Kim, M., Kwon, J.S., 2021. Thalamic connectivity system across psychiatric disorders: Current status and clinical implications. Biol. Psychiatry Glob. Open Sci. 2, 332–340. 10.1016/j.bpsgos.2021.09.008.

Kandilarova, S., Stoyanov, D., Kostianev, S., Specht, K., 2018. Altered resting state effective connectivity of anterior insula in depression. Front. Psychiatry. 9, 83. 10.3389/fpsyt.2018.00083

Koenigs, M., Huey, E. D., Calamia, M., Raymont, V., Tranel, D., Grafman, J., 2008. Distinct Regions of Prefrontal Cortex Mediate Resistance and Vulnerability to Depression. J Neurosci, 28(47), 12341–12348. 10.1523/JNEUROSCI.2324-08.2008.

Leech, R., Sharp, D.J., 2014. The role of the posterior cingulate cortex in cognition and disease. Brain. 137, 12–32. 10.1093/brain/awt162.

LeWinn, K.Z., Connolly, C.G., Wu, J., Drahos, M., Hoeft, F., Ho, T.C., Simmons, A.N., Yang, T.T., 2014. White matter correlates of adolescent depression: structural evidence for frontolimbic disconnectivity. J. Am. Acad. Child Adolesc. Psychiatry. 53, 899–909, 909.e1–7. 10.1016/j.jaac.2014.04.021.

Menon, V., Uddin, L.Q., 2010. Saliency, switching, attention and control: a network model of insula function. Brain Struct. Funct. 214, 655–667. 10.1007/s00429-010-0262-0.

Mincic, A.M., 2015. Neuroanatomical correlates of negative emotionality-related traits: A systematic review and meta-analysis. Neuropsychologia. 77, 97–118. 10.1016/j.neuropsychologia.2015.08.007.

Nemeroff, C.B., 2020. The state of our understanding of the pathophysiology and optimal treatment of depression: Glass half full or half empty? Am. J. Psychiatry. 177, 671–685. 10.1176/appi.ajp.2020.20060845.

Nierenberg, A.A., DeCecco, L.M., 2001. Definitions of antidepressant treatment response, remission, nonresponse, partial response, and other relevant outcomes: a focus on treatment-resistant depression. J. Clin. Psychiatry. 62, 5–9. PMID: 11480882.

Pessoa, L., Adolphs, R., 2010. Emotion processing and the amygdala: from a ‘low road’ to ‘many roads’ of evaluating biological significance. Nat. Rev. Neurosci. 11, 773–783. 10.1038/nrn2920.

Philip, N.S., Arulpragasam, A.R., 2023. Reaching for the unreachable: low intensity focused ultrasound for non-invasive deep brain stimulation. Neuropsychopharmacology. 48, 251–252. 10.1038/s41386-022-01386-2.

Rolls, E.T., 2019. The cingulate cortex and limbic systems for emotion, action, and memory. Brain Struct. Funct. 224, 3001–3018. 10.1007/s00429-019-01945-2.

Rolls, E.T., Cheng, W., Feng, J., 2020. The orbitofrontal cortex: reward, emotion and depression. Brain Commun. 2, fcaa196. 10.1093/braincomms/fcaa196.

Runia, N., Yücel, D.E., Lok, A., de Jong, K., Denys, D.A.J.P., van Wingen, G.A., Bergfeld, I.O., 2022. The neurobiology of treatment-resistant depression: A systematic review of neuroimaging studies. Neurosci. Biobehav. Rev. 132, 433–448. 10.1016/j.neubiorev.2021.12.008.

Sanchez, S.M., Tsuchiyagaito, A., Kuplicki, R., Park, H., Postolski, I., Rohan, M., Paulus, M.P., Guinjoan, S.M., 2023. Repetitive negative thinking-specific and -nonspecific white matter tracts engaged by historical psychosurgical targets for depression. Biol. Psychiatry. 94, 661–671. 10.1016/j.biopsych.2023.03.012.

Sliz, D., Hayley, S., 2012. Major depressive disorder and alterations in insular cortical activity: A review of current functional magnetic imaging research. Front. Hum. Neurosci. 6, 323. 10.3389/fnhum.2012.00323.

Sobstyl M, Kupryjaniuk A, Prokopienko M, Rylski M. Subcallosal Cingulate Cortex Deep Brain Stimulation for Treatment-Resistant Depression: A Systematic Review. Front Neurol. 2022 Apr 1;13:780481. doi: 10.3389/fneur.2022.780481. PMID: 35432155; PMCID: PMC9012165.

Ten Doesschate, F., Bruin, W., Zeidman, P., Abbott, C.C., Argyelan, M., Dols, A., Emsell, L., van Eijndhoven, P.F.P., van Exel, E., Mulders, P.C.R., Narr, K., Tendolkar, I., Rhebergen, D., Sienaert, P., Vandenbulcke, M., Verdijk, J., van Verseveld, M., Bartsch, H., Oltedal, L., van Waarde, J.A., van Wingen, G.A., 2023. Effective resting-state connectivity in severe unipolar depression before and after electroconvulsive therapy. Brain Stimul. 16, 1128–1134. 10.1016/j.brs.2023.07.054.

Tsolaki, E., Narr, K.L., Espinoza, R., Wade, B., Hellemann, G., Kubicki, A., Vasavada, M., Njau, S., Pouratian, N., 2021. Subcallosal cingulate structural connectivity differs in responders and nonresponders to electroconvulsive therapy. Biol. Psychiatry Cogn. Neurosci. Neuroimaging. 6, 10–19. 10.1016/j.bpsc.2020.05.010.

Tsuchiyagaito, A., Misaki, M., Cochran, G., Philip, N.S., Paulus, M.P., Guinjoan, S.M., 2023. Thalamo-cortical circuits associated with trait- and state-repetitive negative thinking in major depressive disorder. J. Psychiatr. Res. 168, 184–192. 10.1016/j.jpsychires.2023.10.058.

Tura, A., Goya-Maldonado, R., 2023. Brain connectivity in major depressive disorder: a precision component of treatment modalities? Transl. Psychiatry. 13, 196. 10.1038/s41398-023-02499-y.

Uddin, L.Q., Nomi, J.S., Hébert-Seropian, B., Ghaziri, J., Boucher, O., 2017. Structure and function of the human insula. J. Clin. Neurophysiol. 34, 300–306. 10.1097/WNP.0000000000000377.

Velasco, F., Velasco, M., Jiménez, F., Velasco, A.L., Salin-Pascual, R., 2005. Neurobiological background for performing surgical intervention in the inferior thalamic peduncle for treatment of major depression disorders. Neurosurgery, 57(3), 439–448. 10.1227/01.neu.0000172172.51818.51.

Yang, C., Xiao, K., Ao, Y., Cui, Q., Jing, X., Wang, Y., 2023. The thalamus is the causal hub of intervention in patients with major depressive disorder: Evidence from the Granger causality analysis. Neuroimage Clin., 37, 103295. 10.1016/j.nicl.2022.103295.

Yeo, B.T., Krienen, F.M., Sepulcre, J., Sabuncu, M.R., Lashkari, D., Hollinshead, M., Roffman, J.L., Smoller, J.W., Zöllei, L., Polimeni, J.R., Fischl, B., Liu, H., Buckner, R.L., 2011. The organization of the human cerebral cortex estimated by intrinsic functional connectivity. J. Neurophysiol. 106, 1125–1165. 10.1152/jn.00338.2011.

Zhao, H., Turel, O., Bechara, A., & He, Q., 2023. How distinct functional insular subdivisions mediate interacting neurocognitive systems. Cereb Cortex, 33(5), 1739–1751. 10.1093/cercor/bhac169.

