## Supplementary Material_ Samman et al 2025 for "Structural Connectivity Correlates of Response to Electroconvulsive Therapy in Treatment-Resistant Depression"

Structural Connectivity Correlates of Response to Electroconvulsive Therapy in Treatment-Resistant Depression

Supplementary Material
Participants
Patients with TRD eligible to receive ECT, were recruited if they met the following inclusion criteria a) a clinical diagnosis of MDD according to DSM-5 as independently assessed by two specialists in psychiatry, b) age between 18 and 70 years, c) stable medication for at least two weeks, and d) treatment resistance defined as two failed antidepressant trials of sufficient duration (4 weeks) and dose, and one augmentation attempt with atypical antipsychotics, an antidepressant of a different chemical group, lithium, or a thyroid hormone. The exclusion criteria for all participants were major neurocognitive disorder, a diagnosis of neurodegenerative conditions like Alzheimer’s Disease and Idiopathic Parkinson’s Disease, a history of at least moderate Traumatic Brain Injury, and a diagnosis of substance use disorder for the last 6 months except nicotine and caffeine.

ECT treatment 
The ECT protocol involved 12 sessions administered according to the following pattern: three weekly sessions for 2 weeks, two weekly sessions for 2 weeks and weekly for 2 weeks. Patients were under general anesthesia with intravenous propofol (2 mg/kg) and muscle relaxation with intravenous succinylcholine (1 mg/kg). A SpECTrum 5000Q (MECTA) device was used to provide pulses of square wave and ultra-short duration with a bilateral frontal electrode placement. A generalized tonic-clonic seizure lasting at least 15 seconds was considered an appropriate response. The seizure duration was assessed both clinically (observing tonic-clonic movements in the left foot, which was not exposed to succinylcholine, by insufflating a sphygmomanometer above 250 mmHg before the administration of succinylcholine, until the end of the seizure) and in a simultaneous 2-lead electroencephalogram recording. Vital signs were continuously monitored by an anesthesiologist. 

Exclusions
Two patients with treatment resistant depression who received ECT (1 female) were excluded from the study because of catatonia symptoms impeding the assessment of cognitive status and provision of informed consent, and other four patients dropped out of treatment during the ECT sessions. One control subject was found to have a HAM-D score of 10. Given that one of the exclusion criteria was the presence of significant depressive symptoms, defined as a HAM-D score of 10 or higher, this individual was subsequently excluded from the HC group.


Image Acquisition 
The participants were scanned on a 3T GE Discovery 750 MRI system (Fleni) within one week prior to the first ECT session. Using a diffusion weighted imaging (DWI) sequence, 47 axial slices were obtained (TR = 4500 ms, TE = 63.1 ms, matrix size = 256x256, voxel size = 0.94x0.94x3mm3) along 60 diffusion directions (b = 1000 s/mm2) and 7 images without diffusion weighting (b0) for each individual. In addition, a high resolution T1 3D fast SPGR-IR image was acquired (TR = 8.272 ms, TE = 3.26 ms, TI = 450); correction (ASSET) with acceleration factor = 2; acquisition matrix = 256x256; FOV = 24cm; slice thickness = 1.1 mm, spacing between slices = 1.1 mm, and 146 axial slices.

Image processing 
Tools from the FSL library (http://www.fmrib.ox.ac.uk/fsl) were used to pre-process the diffusion data and for probabilistic tractography analysis. For each subject, diffusion data were first skull-stripped using the FSL Brain Extraction Tool (BET; Behrens et al., 2003). Eddy current correction for motion and distortion was then performed, including affine registrations of each volume in the diffusion dataset to the initial reference B0 volume. 
Before performing tractography, all diffusion data were processed using BEDPOSTX (from FMRIB's diffusion toolbox, FDT v2.0, http://fsl.fmrib.ox.ac.uk/fsl/fslwiki/FDT) to build up distributions on diffusion parameters at each voxel. This model uses Bayesian techniques to estimate a probability distribution function (PDF) on the principal fiber direction at each voxel, accounting for the possibility of crossing fibers within each voxel. The probability distributions were computed for two fiber directions at each voxel using a multiplicative factor of 1 (Behrens et al., 2007) and 1000 iterations before sampling. Finally, two-step transformation (FLIRT and FNIRT) was used to register scans at native space to the MNI 152 standard space (Jenkinson & Smith, 2001; Zhu et al., 2023) . 

In order to perform the tractography in native space, seed region masks were transformed into each individual’s T1 and diffusion imaging space using FLIRT-FNIRT linear and non-linear transformations. To compute the structural connectivity we used the PROBTRACKX function in FDT toolbox, including 5000 streamlines emerging per seed voxel, according the following termination criteria: the streamline leaves the B0 mask; the streamline loops back on itself or the curvature of the next step was too high compared with the previous one (the curvature threshold was set in 0.2).

References

Behrens, T.E., Woolrich, M.W., Jenkinson, M., Johansen-Berg, H., Nunes, R.G., Clare, S., Matthews, P.M., Brady, J.M., Smith, S.M., 2003. Characterization and propagation of uncertainty in diffusion-weighted MR imaging. Magn. Reson. Med. 50(5), 1077–1088. https://doi.org/10.1002/mrm.10609.

Behrens, T.E., Berg, H.J., Jbabdi, S., Rushworth, M.F., Woolrich, M.W., 2007. Probabilistic diffusion tractography with multiple fibre orientations: What can we gain? Neuroimage 34(1), 144–155. https://doi.org/10.1016/j.neuroimage.2006.09.018.

Jenkinson, M., Smith, S., 2001. A global optimisation method for robust affine registration of brain images. Med. Image Anal. 5(2), 143–156. https://doi.org/10.1016/s1361-8415(01)00036-6.

Zhu, Z., Han, J., Zhu, H., Cai, C., Feng, C., Guo, X., Ying, Y., Jiang, H., Zheng, Z., Zhang, J., Zhu, J., Wu, H., 2023. Individualized targeting is warranted in subcallosal cingulate gyrus deep brain stimulation for treatment-resistant depression: A tractography analysis. Hum. Brain Mapp. 44(11), 4200–4210. https://doi.org/10.1002/hbm.26339.
